# Immunometabolic hijacking of immune cells by a *Pseudomonas aeruginosa* quorum-sensing signal

**DOI:** 10.1101/2021.10.03.462785

**Authors:** Arunava Bandyopadhaya, Vijay K Singh, Arijit Chakraborty, A. Aria Tzika, Laurence G Rahme

## Abstract

Macrophages utilize metabolic pathways to generate energy and metabolites that may be vulnerable to pathogen hijacking to favor pathogen survival and persistence. It is unclear how bacterial pathogens alter metabolic pathways in immune cells for their benefit and persistence in the infected host. We have shown that the *Pseudomonas aeruginosa* quorum sensing (QS) signal molecule 2-aminoacetophenone (2-AA) allows pathogen persistence in host tissues by triggering host tolerization via histone deacetylase (HDAC)1-mediated epigenetic reprogramming. Here, we provide strong evidence that 2-AA-meditated persistence is linked to specific metabolic pathway alterations that reduce energy availability and biosynthetic macromolecules involved in host immune responses. 2-AA promotes a Warburg-like metabolic reprogramming effect, thereby increasing levels of lactate, which repressed inflammatory signaling in macrophages. Moreover, it interferes with pyruvate translocation to mitochondria, reducing mitochondrial (mt)-oxidative phosphorylation (OXPHOS) due to down-regulation of estrogen-regulated receptor (ERR)α and mitochondrial pyruvate carrier *(MPC)-1*. This metabolic reprogramming dampened energy production, reduced the acetyl-CoA pool, and generated an anti-inflammatory milieu that favors *P. aeruginosa* persistence. These findings provide evidence of first-in-class metabolic reprogramming in immune cells mediated by a QS signaling molecule. The specific metabolic programs affected provide insights that may guide the design and development of therapeutics and protective interventions against pathogen-induced immunometabolic alterations and persistence factors.

## Introduction

Host-pathogen interactions are characterized by a continuous antagonistic interplay between pathogen and host (1, 2). *Pseudomonas aeruginosa*, a recalcitrant ESKAPE pathogen, excretes virulence-associated low molecular weight molecules. The synthesis of several of these molecules is regulated by quorum sensing (QS), a cell density-dependent signaling system utilized by bacteria to synchronize their activities (3-5). Some bacterial QS signals impact host immune functions (6). MvfR (a.k.a. PqsR) is a critical QS transcription factor of *P. aeruginosa* that regulates the synthesis of about sixty *P. aeruginosa*-excreted small molecules, including 2-aminoacetophenone (2-AA). 2-AA, which is abundantly produced by *P. aeruginosa* and other pathogens (7), has been shown to promote chronic infection phenotypes via effects on the pathogen itself (8, 9) and its host (1, 10).

In response to a first-time encounter with a pathogen, innate immune cells in host organisms can develop long-term memory and thus undergo reprogramming. Subsequent exposure may enhance or repress immune cell function via processes known as trained immunity and tolerance, respectively (11). Both processes are associated with profound intracellular metabolic and epigenetic reprogramming (1, 11). We have shown that a host’s first exposure to 2-AA leads to reprogramming innate immune cells, whereas a subsequent secondary stimulation with 2-AA reveals immune cell function repression promoted by this molecule. 2-AA modulates the host inflammatory signaling cascade, maintaining chromatin in a “silent” state by increasing Histone DeACetylase (HDAC)1 expression and activity while decreasing Histone AcetylTransferase (HAT) activity (1). The effects of 2-AA on host cells result in regulation of the protein-protein interactions between HAT and cyclic AMP response element-binding protein/HDAC1 and p50/p65 nuclear factor (NF)-κB subunits (1, 12). The epigenetic mechanism and effects mediated by 2-AA are distinct from previously described bacterial molecules that repress inflammatory responses. 2-AA-mediated tolerization permits *P. aeruginosa* to persist in infected tissues (1), an outcome that is distinct from lipopolysaccharide (LPS) tolerization, which instead leads to bacterial clearance (13) and involves different HDACs (14).

Metabolic pathways contribute to histone acetylation and deacetylation modulation processes that mediate transcriptional reprogramming of immune cells during infection (15). Epigenetic regulation can be controlled by intermediate metabolites of metabolic pathways serving as substrates to epigenetic enzymes (e.g., acetyl-CoA, the primary substrate for histone acetyltransferases) (16). Several reports point to the activation of specific metabolic programs in monocytes and macrophages upon exposure to microbial products (17, 18). Intracellular bacterial infections trigger metabolic reprograming in mammalian cells that alter metabolic pathways and levels of metabolites in infection microenvironments (19). In turn, macrophages utilize metabolic pathways to generate energy and metabolites and adapt rapidly to changing environments and stimuli, thereby enabling them to cope with the fluctuating immune-response needs (19).

Current working models of immune-cell metabolic reprogramming are characterized by increased cellular aerobic glycolysis and impaired mitochondrial (mt) oxidative phosphorylation (OXPHOS) (20). This switch from OXPHOS to glycolysis resembles the Warburg effect observed in cancer cells (21) and chronic diseases (22). Training of immune cells is associated with profound intracellular metabolic reprogramming and epigenetic modifications. The fungal cell wall component β-glucan and the Bacillus Calmette-Guérin vaccine are known to bring about posttranslational modifications of histones associated with modulation of the expression of genes involved in glycolytic and OXPHOS metabolism, thus linking metabolic changes to epigenetic reprogramming (11). In the case of LPS-mediated tolerance, HDAC activation is promoted by glycolytic shifts concurrent with OXPHOS suppression in immune cells (23). There is also evidence of a reverse-causal link, whereby impeding the activation of aerobic glycolysis prevents the characteristic chromatin modification pattern and adapted phenotype of trained immunity (24). However, the mechanisms and processes underlying the metabolic changes occurring in immune cells following bacterial infection remain poorly understood.

The purpose of this study is to interrogate mechanistic aspects and processes that contribute to the host metabolic reprogramming and maintenance of host tolerance-mediated pathogen persistence that are conferred by the QS signaling molecule 2-AA. Given that 2-AA mediated tolerization in immune cells is maintained by epigenetic reprograming (1) and our previous *in vivo* findings in skeletal muscle showed that 2-AA dampens ATP synthesis rate, acetyl-CoA biosynthesis, the citric acid cycle (CAC), fatty acid biosynthesis, electron transport, and OXPHOS (1, 25, 26), here, we conducted a series of *in vitro* and *in vivo* experiments to interrogate whether the epigenetic reprogramming previously observed in macrophages is linked to alteration of their cellular metabolism (1, 12). In murine *in vivo* studies, we also assessed bacterial persistence in tolerized and non-tolerized mice to corroborate our immunometabolic findings and our previous findings showing that 2-AA tolerization increased survival of infected mice by 90%, dampened cytokine storm, and led to a persistent high bacteria load which was sustained in infected mice (10, 12). Finally, to further pin down 2-AA as the culprit of the observed immunometabolic changes, we infected macrophages with wild-type *P. aeruginosa* (PA14) and two isogenic mutants, namely an *mvfR* mutant and a *pqsA* mutant, that do not produce 2-AA (9). Understanding the immunometabolic reprogramming mediated by 2-AA may reveal critical insights into the host-pathogen interplay that leads to tolerization and clinical recalcitrant pathogen persistence.

## Results

### 2-AA tolerization promotes glycolysis and compromises OXPHOS in immune cells

The experimental design used to assess lactate, pyruvate, acetyl-CoA, and ATP in 2-AA-tolerized and non-tolerized mouse macrophages (RAW264.7) and human monocytes (THP-1) cells, respectively, are shown in Figures 1A and S1A. Because lactate plays a key role in energy regulation, immune tolerance, and immune memory, we assessed its intracellular and extracellular levels in 2-AA-tolerized and non-tolerized macrophages (Fig. 1B & Fig. S1B). Stimulation of macrophages with 2-AA led to a gradual induction of lactate over 24 h. Maximum levels were reached within 1 h and maintained for at least 24 h. The high lactate levels remained unchanged in tolerized cells, even following the second 2-AA stimulation. 2-AA non-tolerized and tolerized THP-1 human monocytes exhibited similar patterns in lactate levels (Fig. S2A): an increase in intracellular lactate dehydrogenase (LDH) activity (Fig. 1C), with increased LDH-mediated (reverse) catalysis of pyruvate reduction into lactate over time, in the absence of observable extracellular LDH activity or cytotoxicity (Fig. S2B).

**Figure 1.**
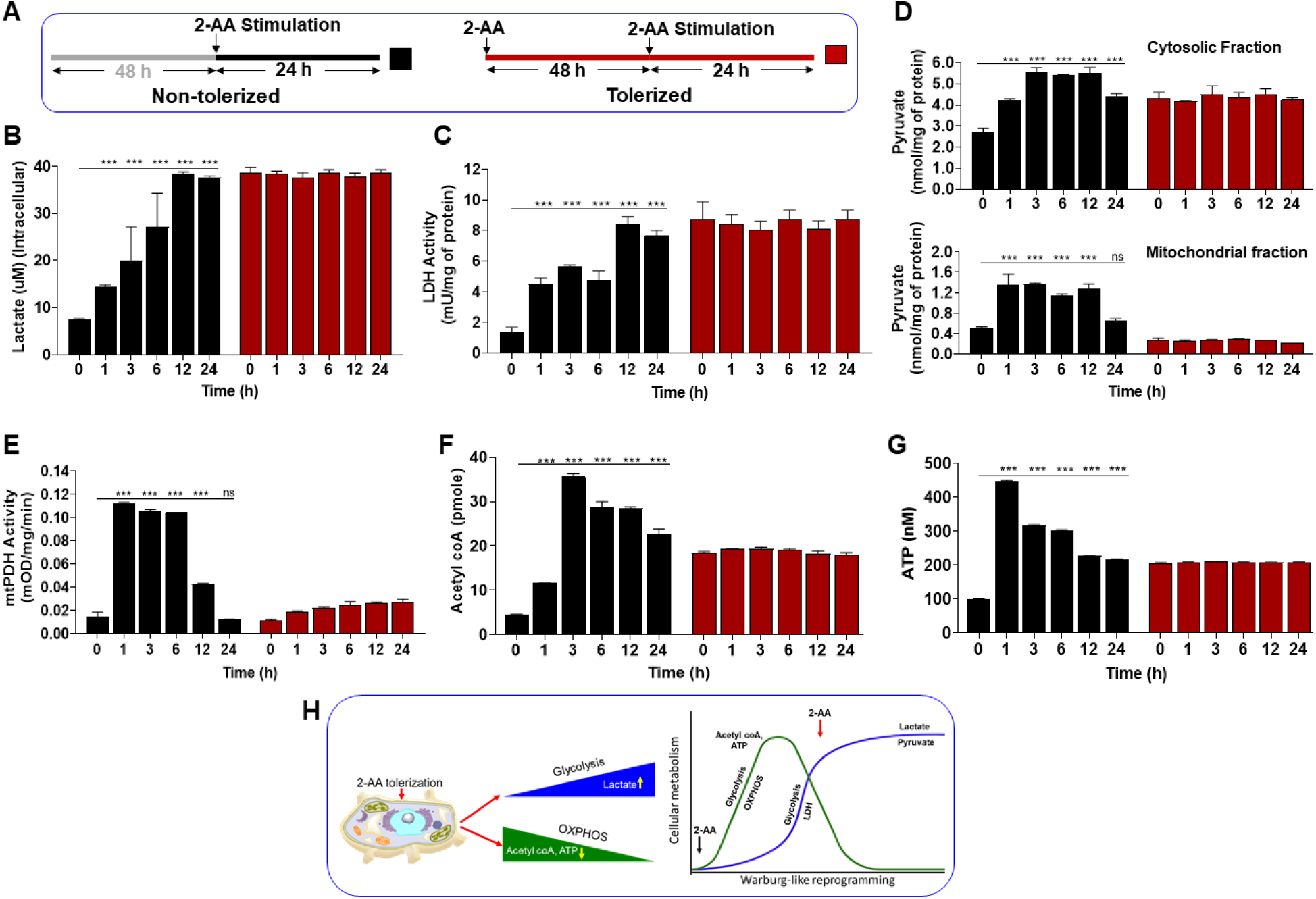
2-AA tolerization leads to decreased metabolites in mouse macrophages. (A) Mouse macrophage RAW264.7 cells were tolerized with 2-AA for indicated time periods (up to 48 h) or left untreated. (B) Intracellular lactate, (C) intracellular LDH activity, (D) cytosol and mitochondria pyruvate content, (E) mt-PDH activity, (F) acetyl-CoA, and (G) ATP were measured in non-tolerized and 2-AA-tolerized RAW264.7 cells following 2-AA stimulation (2-AA Sti) at indicated time points. Means ± SDs are shown (n = 3); \*\*\**p* < 0.001 and ns indicates no significant difference for one-way ANOVA. (H) Schematic representation of 2-AA mediated reprogramming of metabolism. Following cell stimulation, increased metabolism generated acetyl-CoA and ATP; subsequent intracellular lactate and pyruvate accumulation prevented OXPHOS. Upon subsequent stimulation, 2-AA induced lactate and pyruvate accumulation promoted a glycolytic state via Warburg-like reprogramming.

To explain the lactate accumulation observed in 2-AA tolerized cells, we assessed pyruvate levels in cytosol and mitochondria fractions of mouse macrophages (Fig. 1D) and human monocytes (Fig. S1C). Compared to non-tolerized control cells (RAW264.7 and THP-1, respectively), the pyruvate levels were decreased in the mitochondria fraction while being highly accumulated in the cytosolic fraction of 2-AA tolerized cells. In contrast, non-tolerized cells exhibited pyruvate increases in both fractions over time (Fig. 1D & S1C).

In mitochondria, pyruvate is converted to acetyl-CoA (Fig. 2A), the primary substrate for histone acetyltransferases, by mt-pyruvate dehydrogenase (PDH). Assessment of mt-PDH activity (Fig. 1E) in non-tolerized cells demonstrated an increase in mt-PDH activity within 1 h of tolerization, followed by a decline by 12 h. Meanwhile, in 2-AA tolerized cells, mt-PDH activity was sustained at basal levels.

**Figure 2.**
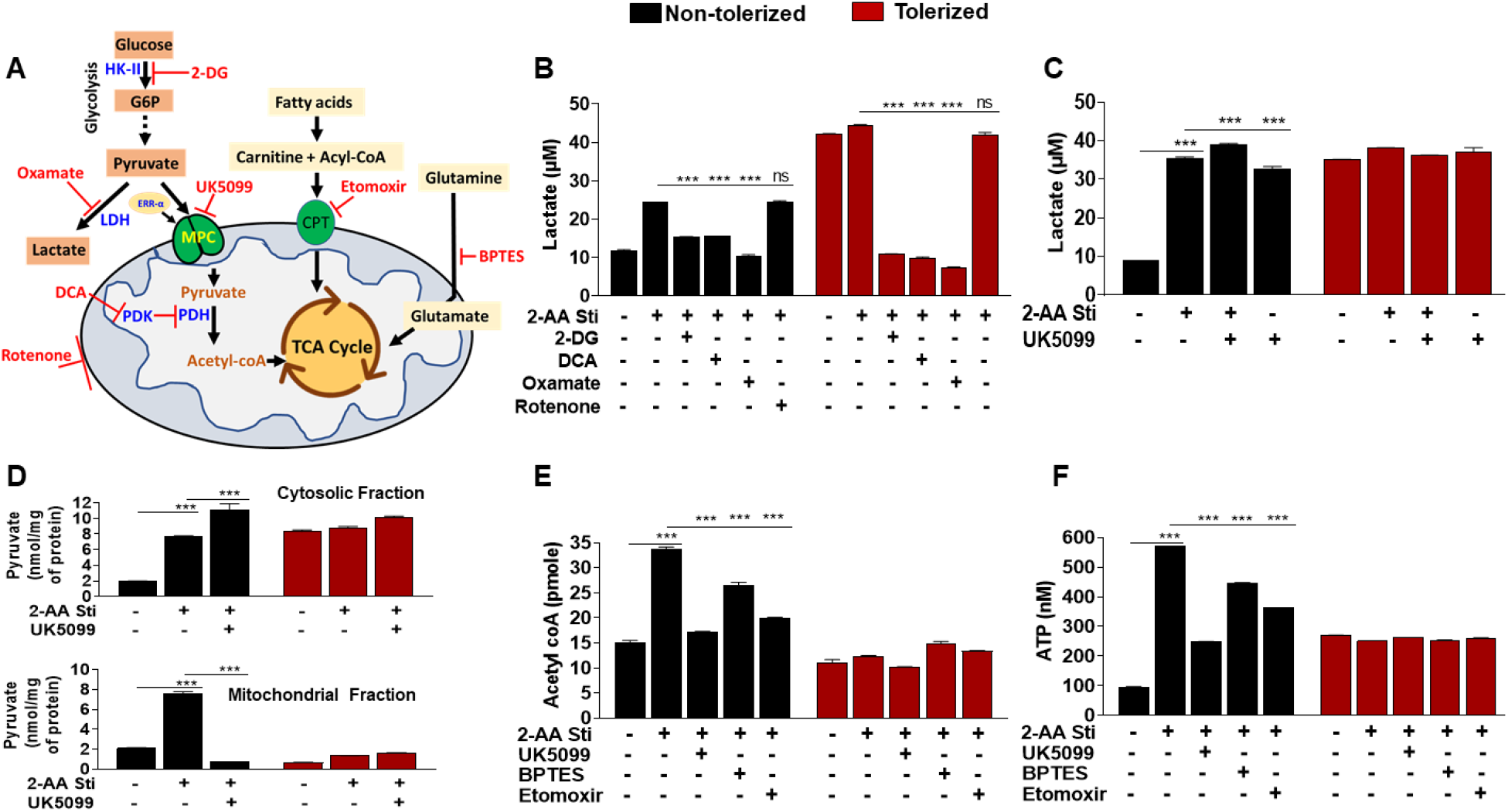
2-AA tolerization promotes glycolytic fluxes and reduces TCA fluxes, leading to increased lactate production in mouse macrophages. (A) Inhibitors for key glycolysis and mt-OXPHOS pathways. (B) Intracellular lactate levels in non-tolerized and 2-AA-tolerized RAW264.7 cells following 2-AA stimulation (2-AA Sti) ± 2-DG, ± DCA, and ± oxamate, ± rotenone for 24 h. (C) Cellular lactate in non-tolerized and 2-AA tolerized RAW264.7 cells following 2-AA stimulation ± UK5099 24 h. (D) Cellular and mt-pyruvate levels in non-tolerized and 2-AA tolerized RAW264.7 cells following 2-AA stimulation ± UK5099 3 h. (E-F) Levels of cellular ATP (E) and acetyl-CoA (F) in non-tolerized and 2-AA-tolerized RAW264.7 cells following 2-AA stimulation ± UK5099, ± BPTES, and ± etomoxir for 3 h. Means ± SDs are shown (n = 3); \*\*\**p* < 0.001 and ns indicates no significant difference for one-way ANOVA.

2-AA non-tolerized mouse macrophages (Fig. 1F & G) and human monocytes (Fig. S1D & E) exhibited induced acetyl-CoA and ATP generation within 1 h. In contrast, 2-AA tolerized cells exhibited low and sustained levels of acetyl-CoA and ATP and showed no change in ATP generation compared to non-tolerized cells (Figs. 1G & S1E). Given that decreased pyruvate levels and PDH activity in mitochondria can impair acetyl-CoA and subsequent ATP production, our results suggest that 2-AA tolerization promoted a glycolytic shift characterized by lactate accumulation and compromised OXPHOS via repression of acetyl-CoA and ATP generation in macrophages and human monocytes (Fig. 1H).

### 2-AA tolerization induces glycolytic fluxes in macrophages

Lactate levels are determined by the balance between glycolysis and mt-metabolism pathways (ref). We tested if the glycolytic shift observed is due to increased glycolytic fluxes in 2-AA tolerized cells. Lactate production is induced by glucose; the non-metabolizable glucose analog 2-deoxy-d-glucose (2-DG) (27) decreased lactate production in 2-AA stimulated and tolerized cells (Figs. 2A & B). Sodium dichloroacetate (DCA) and oxamate were used to inhibit lactate production by modulating the activities of PDH (28) and LDH (29), respectively (Fig. 2B). These two compounds decreased the intracellular levels of lactate in 2-AA tolerized and non-tolerized cells (Fig. 2B). Rotenone, an inhibitor of the mitochondrial respiratory chain complex I (30), which drives cells towards glycolysis, did not alter intracellular lactate levels in 2-AA tolerized or non-tolerized cells (Fig. 2B). These findings suggest that 2-AA-tolerization increases glycolytic flux with consequently increased lactate levels that, in turn, could be a result of decreased oxidative metabolism in immune cells.

### 2-AA tolerization modulates mt-pyruvate flux and -OXPHOS by regulating pyruvate carrier (MPC) functions

To interrogate the relevance of mt-OXPHOS and relative utilization of glycolysis in 2-AA tolerized cells, we used UK5099, which restricts pyruvate entry in mitochondria by inhibiting MPC1 function, (31) BPTES (inhibitor of glutamine-driven OXPHOS) (32), and etomoxir (inhibitor of fatty acid oxidation) (33) (Fig. 2A). UK5099 addition did not change accumulated lactate levels in 2-AA tolerized or non-tolerized cells, but it did induce lactate accumulation in non-tolerized unstimulated cells (Fig. 2C). UK5099 reduced pyruvate levels in mitochondria fractions, but not in cytosol fractions of 2-AA stimulated cells (Fig. 2D), indicating that 2-AA prevents pyruvate entry into mitochondria by interfering with MPC functions. The addition of UK5099, BPTES, and etomoxir to 2-AA stimulated cell cultures decreased the generation of acetyl-CoA and ATP (*p* < 0.001 vs. no inhibitor controls) (Fig. 2E-F). These inhibitors did not alter levels of acetyl-CoA or ATP in 2-AA tolerized cells (Fig. 2E-F). Together these data show that inhibiting OXPHOS in 2-AA stimulated cells mimics 2-AA tolerization and is sufficient to suppress acetyl-CoA and ATP generation while increasing aerobic glycolysis by modulating MPC functions.

### Down-regulation of estrogen-related receptor alpha (ERR-α) and MPC1 promotes glycolysis state in the 2-AA tolerized cells

The nuclear receptor ERR-α regulates lactate utilization by down-regulating LDH activity and upregulating MPC1 expression to facilitate pyruvate entry into mitochondria (Fig. 2A)(34). Our data show that 2-AA-tolerized cells have decreased ERR-α and MPC1 protein levels together with increased LDH expression (Figs. 3A and S3A, B & F). Down-regulation of ERR-α resulted in modest reductions in both total PDH and phosphorylated PDH levels in 2-AA tolerized cells (Figs. 3A and S3C & D), consistent with PDH being an ERR-α target gene (34). Therefore, 2-AA mediated down-regulation of MPC1 could be mediated by ERR-α down-regulation, impairing pyruvate transport into mitochondria and triggering glycolysis, a feature of the Warburg effect. Moreover, levels of key regulatory proteins of glycolysis [glucose transporter 1 (GLUT1) and hexokinase II (HK2)] and LDH were upregulated in 2-AA-tolerized cells (Figs. 3A and S3E-G). UK5099 added to inhibit pyruvate flux into mitochondria resulted in increased GLUT1, HK-II, and LDH protein expression in 2-AA stimulated and 2-AA tolerized cells (Fig. 3B-E). These results indicate that the ERR-α and MPC-1 down-regulation can induce a glycolytic shift by enabling lactate accumulation and dampening acetyl-CoA/ATP generation by limiting the pyruvate source for mt-OXPHOS in 2-AA tolerized cells.

**Figure 3.**
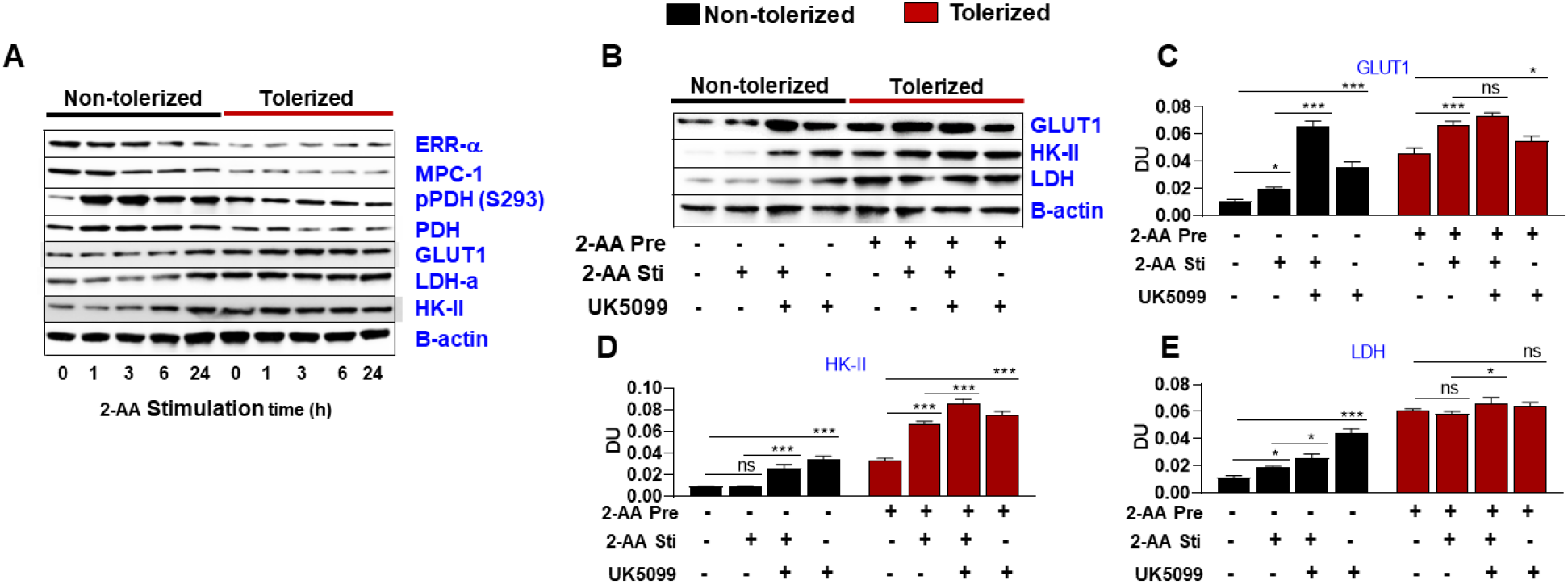
2-AA tolerization modulates the regulator of glycolysis and CAC in macrophages. (A) Representative immunoblot of ERR-α, MPC-1, pPDH, PDH, GLUT1, LDH-α, and HK-II protein levels in non-tolerized and 2-AA-tolerized RAW264.7 cells following 2-AA stimulation (2-AA Sti). (B) Effects of MPC inhibition with UK5099 on GLUT1, HK-II, LDH in non-tolerized and 2-AA-tolerized RAW264.7 cells following 24 h 2-AA stimulation. (C-E) Histograms showing relative expression of (C) GLUT1, (D) HK-II, and (E) LDH in non-tolerized and 2-AA-tolerized RAW264.7 cells following 24 h 2-AA stimulation. β-actin was used as a control. The blots are representative of three independent experiments. Means ± SDs are shown (n = 3); \**p* < 0.05, \*\*\**p* < 0.001, and ns indicates no significant difference for one-way ANOVA.

### HDAC1 knockdown (KD) restores cellular respiration partially in 2-AA tolerized cells

Given that 2-AA mediated tolerization reduces acetyl-CoA and ATP pools, and the concentrations of these molecules affect histone acetylation (1, 12), we examined how HDAC1 impacts glycolytic shifts and dampens OXPHOS in 2-AA tolerized cells. To verify whether 2-AA influences key metabolites, we first compared cellular metabolite levels in control RAW264.7 and HDAC1-KD cells. Compared to tolerized control mouse macrophage cells subjected to 2-AA stimulation, levels of lactate (Fig. 4A), mt-pyruvate (Fig. 4B), acetyl-CoA (Fig. 4C), and ATP (Fig. 4D) were restored in 2-AA-tolerized HDAC1-KD mouse macrophages. These data indicate that HDAC1 may influence glycolytic shifting and reduce acetyl-CoA and energy production in 2-AA tolerized cells by compromising mtOXPHOS.

**Figure 4:**
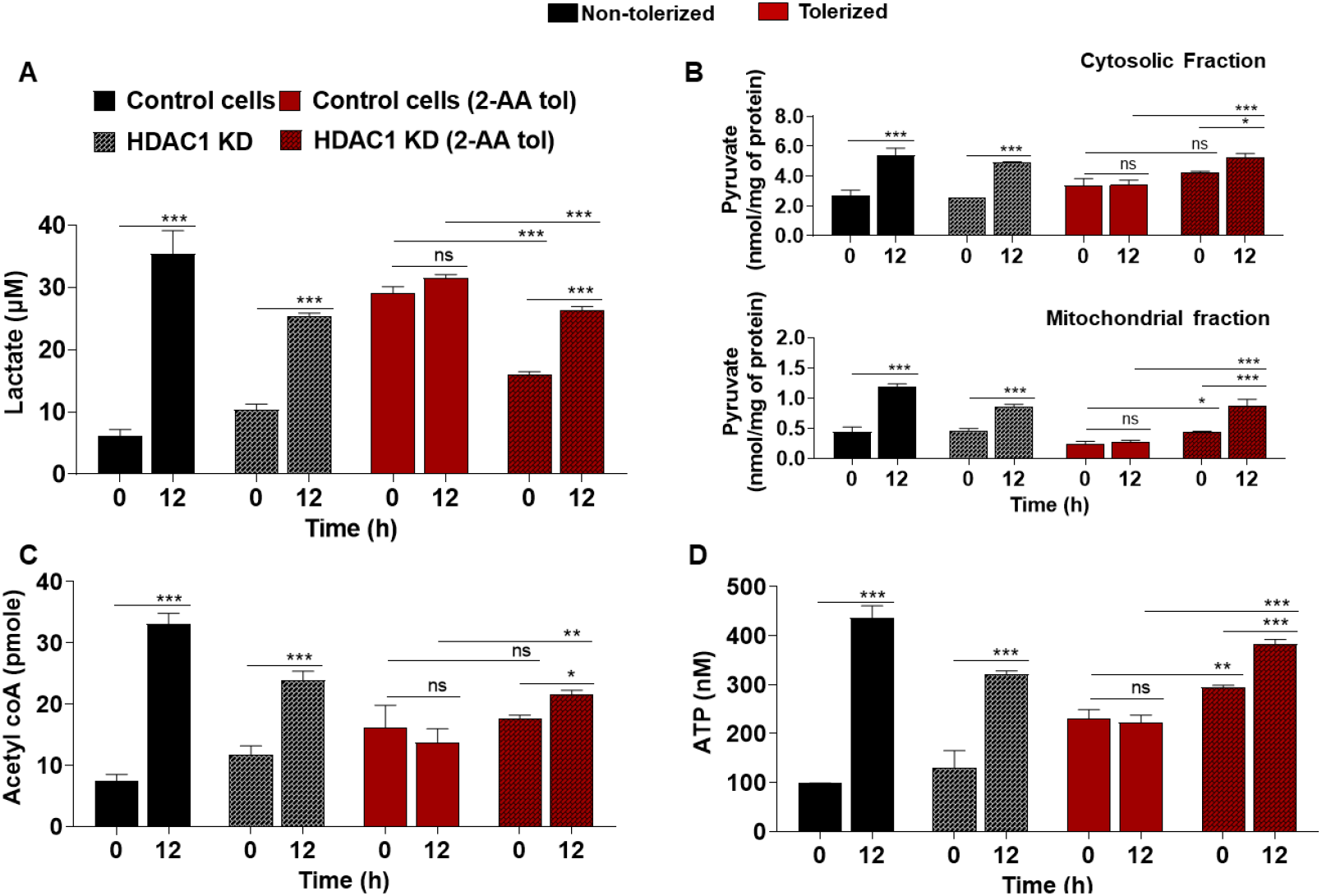
HDAC1 KD restores metabolites in mouse macrophages. (A) Intracellular lactate, (B) cytosolic- and mt-pyruvate, (C) acetyl-CoA content, and (D) cellular ATP in non-tolerized and 2-AA-tolerized wild-type and HDAC1-KD RAW264.7 cells following 2-AA stimulation (2-AA Sti). Means ± SDs are shown (n = 3); \**p* < 0.05, \*\**p* < 0.01, \*\*\**p* < 0.001, and ns indicates no significant difference for one-way ANOVA.

### Metabolic shifts regulate innate immune responses in the 2-AA tolerized cells

Fluctuations in acetyl-CoA and ATP concentrations are translated into dynamic HAT/HDAC activity levels and ultimately affect transcriptional regulation (16). Previously, we have shown that 2-AA tolerization dampens innate immune cell signaling and inflammation *in vivo* and *in vitro* by regulating epigenetics functions in macrophages (1, 10, 12). Thus, here, to determine if the altered acetyl-CoA pool and energy expenditure impact innate immune responses, we assessed HDAC1 activity, TNF-α secretion, histone 3 acetylated at lysine in position 18 (H3K18ac), and NF-κBp65 levels in tolerized and non-tolerized macrophages in the presence or absence of UK5099, BPTES, and etomoxir. HDAC activity was increased in the presence of UK5099 in non-tolerized and 2-AA tolerized cells (Fig. 5A), whereas H3K18ac (Fig. 5B), TNF-α (Fig. 5C), and acetylated NF-κBp65 (Fig. 5D) levels were significantly reduced in non-tolerized cells in the presence of UK5099, BPTES, or etomoxir. These data suggest that compromised OXPHOS can impact histone acetylation, p65 activation, and TNF-α secretion in 2-AA tolerized cells.

**Figure 5.**
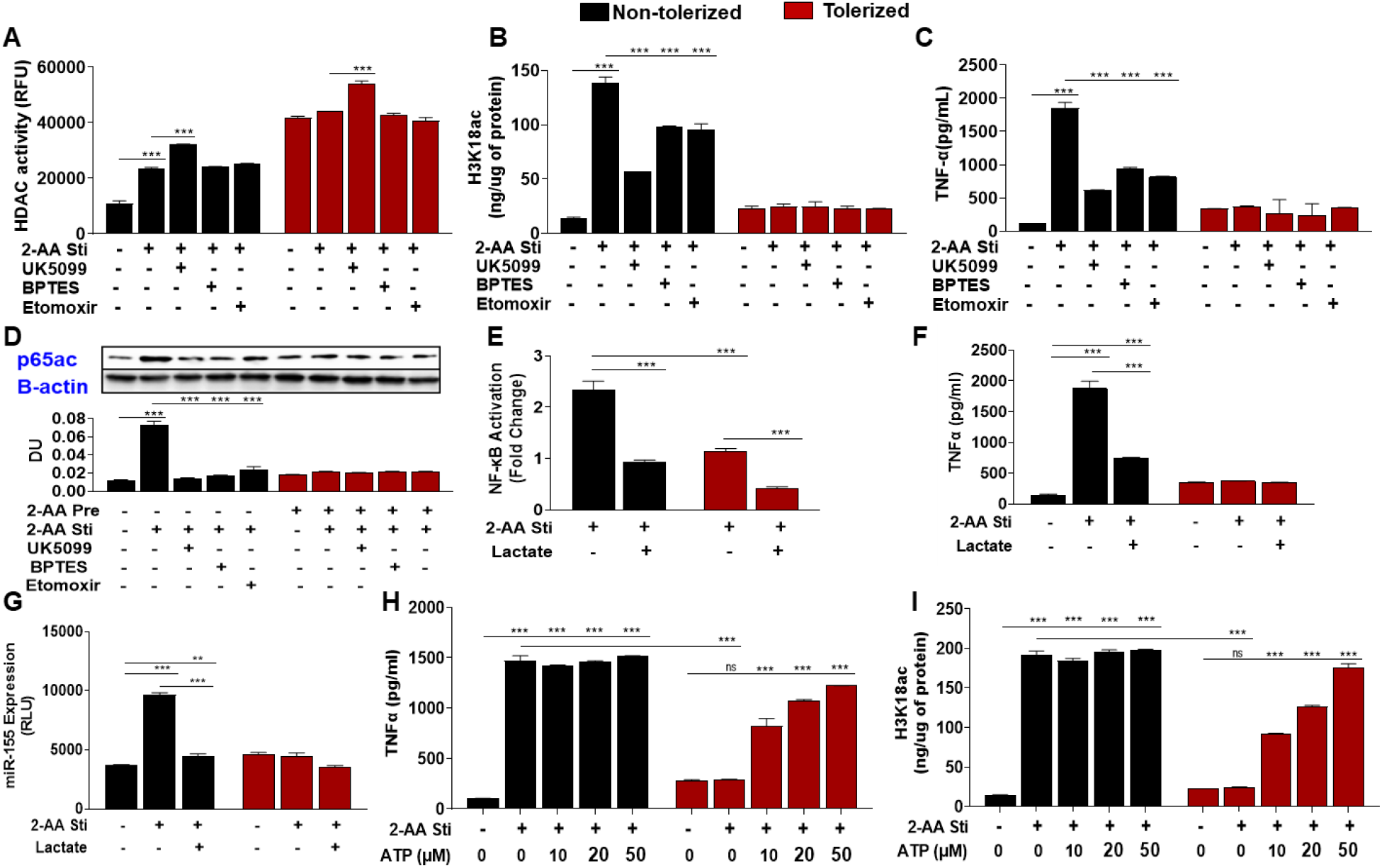
2-AA regulates innate immune function in mouse macrophages by altering metabolite mediated epigenetic regulation. (A) HDAC activity, (B) global H3K18ac protein, and (C) TNF-α levels in non-tolerized (no Pre) and 2-AA-tolerized (2-AA Pre) RAW264.7 cells following 2-AA stimulation (2-AA Sti) ± U5099, ± BPTES, and ±etomoxir for 3 h. (D) Representative immunoblot of NF-κB p65 acetylation was measured in non-tolerized and 2-AA-tolerized RAW264.7 cells following 2-AA stimulation ± U5099, ± BPTES, and ± etomoxir for 1 h. Histograms show the relative expression of proteins; data are representative of three independent experiments. (E-G) NF-κB activity at 2 h (E), TNF-α level at 6 h (F), and miR155 expression at 2 h (G) were analyzed in non-tolerized and 2-AA-tolerized RAW264.7 cells Following 2-AA stimulation ±lactate supplementation. (H-I) TNF-α levels at 6 h (H) and global H3K18ac levels at 1 h (I) were analyzed in non-tolerized and 2-AA-tolerized RAW264.7 cells following 2-AA stimulation ± ATP supplementation. Means ± SDs are shown (n = 3); \*\**p* < 0.01, \*\*\**p* < 0.001, and ns indicates no significant difference for one-way ANOVA. DU, densitometric unit.

### 2-AA tolerization dampens inflammation via lactate accumulation

Because 2-AA tolerization dampens the activity and nuclear translocation NF-κB activity (35), as well as NF-κB-regulated transcription and secretion of TNF-α (1, 10), we examined how lactate affects TNF-α and NF-κB activation. In non-tolerized and 2-AA tolerized cells, lactate supplementation repressed NF-κB activation (Fig. 5E) and TNF-α secretion (Fig. 5F) following 2-AA stimulation. Expression of microRNA (miR)155 (Fig. 5G) was induced in non-tolerized but not in the tolerized cells (Fig. 5G). Lactate supplementation repressed induction of miR155 expression in non-tolerized cells (Fig. 5G), indicating that lactate accumulation may contribute to the 2-AA mediated immune suppression observed in 2-AA tolerized immune cells.

### ATP supplementation reverses 2-AA mediated immunomodulation

ATP is crucial for histone acetylation and gene regulation (36). Because 2-AA tolerization reduces ATP generation and transcription of host defense genes through epigenetic changes in histone acetylation marks, we supplemented non-tolerized and 2-AA tolerized cells with increasing concentrations of ATP. ATP supplementation reversed TNF-α secretion (Fig. 5H) and H3K18 acetylation (Fig. 5I) dose-dependence in tolerized cells.

### *In vivo* studies corroborate the 2-AA-mediated immunometabolic reprogramming

Biochemical analysis of murine spleen tissue obtained from tolerized and non-tolerized mice 1 d, 5 d, and 10 d post-infection (dpi) with *P. aeruginosa* (strain PA14) showed significantly elevated lactate levels (Fig. 6A) over time in tolerized animals. At the 10-dpi time point, acetyl-CoA (Fig. 6B) and ATP (Fig. 6C) levels were significantly reduced in the spleens of infected animals, with reductions being more prominent in tolerized mice than in non-tolerized mice. Bacterial levels in infected mice increased robustly within 24 h of infection (Fig. 6D). In tolerized animals, this load was maintained at a relatively steady level. In non-tolerized animals, it decreased by 5 dpi, but retained a substantial bacterial load at 10 dpi. In infected non-tolerized mice, 2-AA was produced entirely endogenously by *P. aeruginosa*. Meanwhile, in tolerized mice, supplemental exogenous 2-AA was added. These *in vivo* data support ours *in vitro* findings showing that 2-AA tolerization alters metabolic functions and promotes a glycolytic shift.

**Figure 6.**
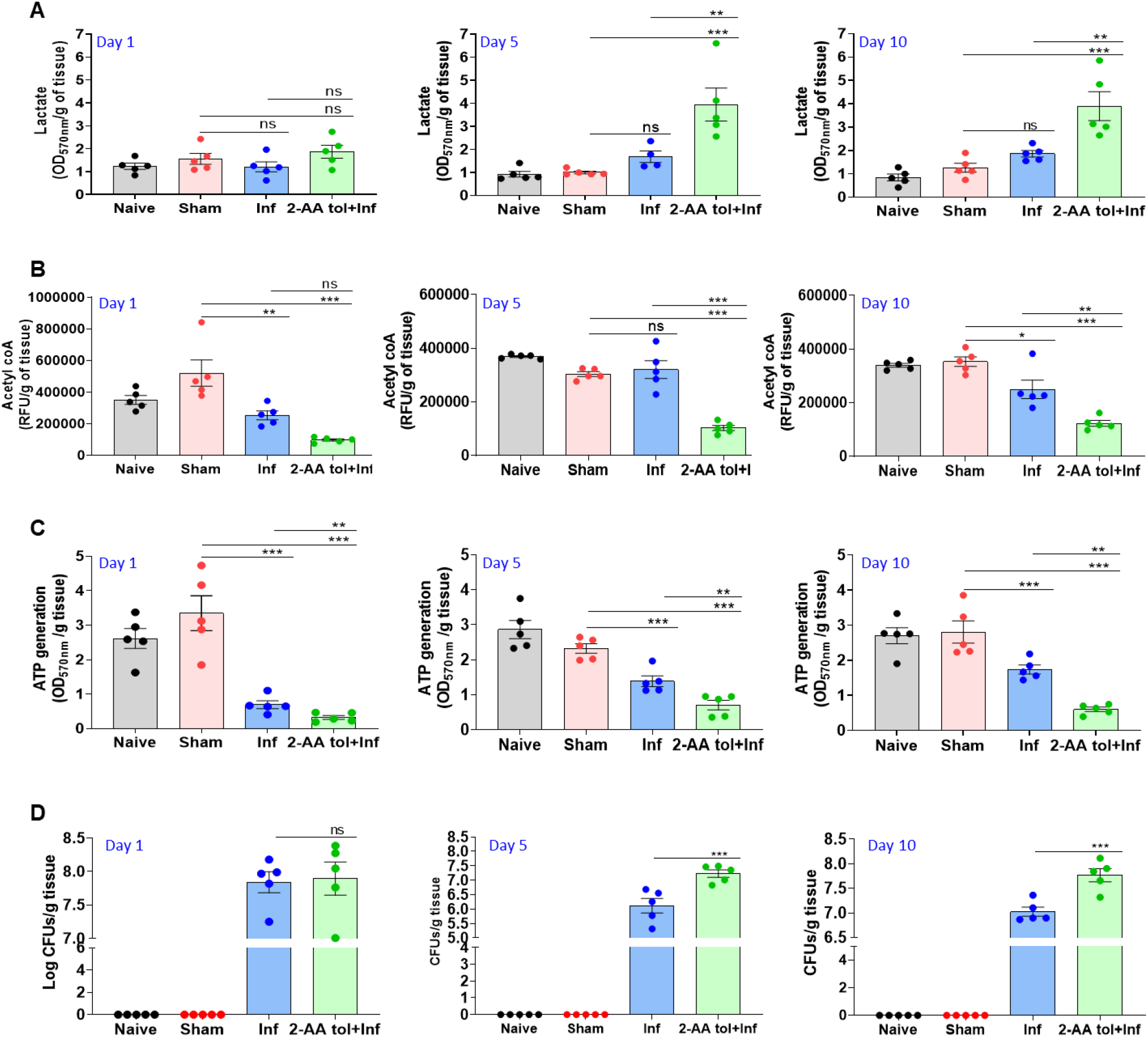
2-AA mediated host tolerance modulates metabolite levels *in vivo* and promotes bacterial persistence. A) Lactate (B) acetyl-CoA, and (C) ATP levels in the spleens of mice treated with 2-AA 4 d prior to undergoing *P. aeruginosa* strain PA14 infection (2-AA tol + Inf). Levels are shown for 1 d, 5 d, and 10 d after infection (Inf). Control mice were not given 2-AA; Sham represents a burn-PBS only group; N = 5 mice/group. Means ± SDs are shown; \**p* < 0.05, \*\**p* < 0.01, \*\*\**p* < 0.001, and ns indicates no significant difference for one-way ANOVA. (D) Bacterial burden (as CFUs) in muscle 1 d, 5 d, and 10 d post-treatment in Inf mice that received 2-AA treatment 4 d prior to Inf. N = 5 mice/group; \*\*\**p* < 0.001 and ns indicates no significant difference for Kruskal-Wallis tests.

Macrophages infected with *mvfR* or *pqsA* mutants of *P. aeruginosa* (strain PA14) exhibited decreased lactate levels (Fig. S4A) together with increased acetyl Co-A (Fig. S4B), ATP (Fig. S4C), and TNF-α (Fig. S4D) levels, as compared to the levels in macrophages infected with wild-type *P. aeruginosa* (PA14). In agreement, macrophages infected with *mvfR* or *pqsA* mutants (Fig. 6E) had reduced HDAC activity levels versus those in macrophages infected with wild-type *P. aeruginosa*. These data support the inference that 2-AA induces immunometabolic reprogramming.

## Discussion

Pathogens hijack host metabolic pathways to avoid clearance and, thus, potentially favor their long-term presence in host tissues (19, 37). Recently, we demonstrated that the QS small molecule 2-AA trains host cells to tolerate a sustained presence of *P. aeruginosa* by way of HDAC1-mediated epigenetic reprogramming of innate immune responses in immune cells (1). The present data show that the 2-AA mediated tolerization of immune cells maintained by epigenetic reprograming is regulated by metabolites and metabolic networks, implicating an interplay between metabolic components and epigenetic regulation in immune cells. This conclusion is supported by the following six findings: (i) 2-AA promoted aerobic glycolysis and repressed mt-metabolism *in vitro* and *in vivo*; (ii) down-regulation of ERR-α and MPC1 compromised OXPHOS and altered ATP and acetyl-CoA levels in 2-AA tolerized cells; (iii) lactate mediated immunomodulation in 2-AA tolerized cells; (iv) ATP supplementation counteracted immunomodulation in 2-AA tolerized cells; (v) increased levels of lactate promoted anti-inflammatory property in 2-AA tolerized cells; and (vi) 2-AA-mediated metabolic shifts observed *in vitro* were recapitulated *in vivo* in *P. aeruginosa* infected mice. These findings provide the first evidence, to our knowledge, of a QS molecule inducing immunometabolic reprogramming in host cells and provide strong support for the notion that 2-AA tolerization involves the reprogramming of glycolytic shifting and the triggering of epigenetic-metabolic interplay in innate immune cells, which, in turn, augments host tolerance to long-term pathogen presence.

Immune cells can detect foreign cells and respond to bacterial infection by altering their metabolic state; meanwhile, some pathogenic bacteria have evolved the ability to evade immune cell activation or adapt to or even benefit from the metabolic shifts that they induce in the immune cells (19). Pathogenic bacteria, including *Chlamydia pneumoniae, Legionella pneumophila*, and *Mycobacterium tuberculosis*, have been shown to shift their host cells towards a Warburg-like metabolism (aerobic glycolysis), enabling them to persist in their hosts (38-40). A recent study revealed that *Salmonella* pathogenesis leads to an accumulation of host glycolytic intermediates that the bacteria use as metabolite sources that support their survival in their hosts (41). *P. aeruginosa-*excreted 2-AA impairs the biosynthesis of CAC intermediates, including acetyl-CoA, in skeletal tissues (25). Cumulatively, our data show that 2-AA reprograms immunometabolism by impairing OXPHOS and altering acetyl-CoA and ATP availabilities. On the other hand, 2-AA shifts host cells’ metabolism towards aerobic glycolysis via accumulation of lactate, a metabolic byproduct of the Warburg effect that represses immune-inflammatory signaling in macrophages. Such metabolic shifts may provide a favorable environmental niche for pathogens that enables them to persist in infected tissues. Future studies should aim to decipher the importance of the host metabolites for bacterial persistence.

ERR-α has been recognized as a master regulator of mt-OXPHOS metabolism (42). Inhibition of ERR-α down-regulates MPC, leading to defective mt-pyruvate uptake and PDH activity, culminating in cellular lactate accumulation and reducing acetyl-CoA and other CAC intermediates (34). If MPC is the primary gatekeeping regulator of carbon sources in the CAC, its down-regulation will result in a bottleneck at the entrance to the CAC. Inhibition of MPC revealed that 2-AA-mediated tolerization compromises acetyl-CoA conversion from pyruvate limits ATP generation and impairs mt-OXPHOS by dampening ERR-α and MPC1. 2-AA is the first QS-regulated small molecule shown to affect ERR-α and MPC1 expression and to be involved in Warburg-like metabolic reprogramming.

An intricate immunometabolic network governs macrophage functions by modifying epigenetic regulation that influences immune cell training during bacterial infection (11). 2-AA tolerizes immune cells by reprograming the host epigenome and dampens inflammation under a high bacterial burden. These effects are distinct from those mediated by β-glucan, Bacillus Calmette-Guérin vaccination (11), or LPS (43), which also promote host protection via epigenetic-metabolic reprogramming. β-glucan stimulation induces a shift from OXPHOS to glycolysis that inhibits the histone deacetylase Sirtuin1 (44). LPS, which promotes endotoxin tolerance in immune cells, has been shown to upregulate glycolysis while suppressing OXPHOS, resulting in increased NAD^+^/NADH ratios, which activate the histone deacetylases Sirtuin 1 and 6, leading to a switch from a proinflammatory state to a more anti-inflammatory state (23). In 2-AA mediated tolerization, the glycolytic shift toward OXPHOS leads to decreased acetyl-CoA and ATP levels, which tends to promote hypoacetylation of proinflammatory loci via maintenance of HDAC1 levels. Acetyl-CoA, which is reduced by 2-AA-mediated tolerization, is a key metabolite that links metabolism with chromatin structure and transcription (16). Thus, acetyl-CoA could underlie the 2-AA-mediated reduction of HAT and maintenance of HDAC1 histone deacetylation, which dampen inflammation (1).

Sustained HDAC1 expression and activity in 2-AA tolerized cells may promote a Warburg-like effect, wherein aerobic glycolysis shifts to mtOXPHOS. Inhibition of HDAC1 results in global disruption of super-enhancers that drive the expression of genes involved in aerobic glycolysis, resulting in enhanced mtOXPHOS (45). Persistent *M. tuberculosis* infection promotes a Warburg-like shift, including upregulation of multiple glycolytic enzymes and glucose uptake transporters, down-regulation of OXPHOS and CAC enzymes (46), and promotion of histone deacetylation by HDACs (47).

Lactate acts in a paracrine manner and contributes to an anti-inflammatory environment by increasing the expression of anti-inflammatory genes (48). A study conducted in human monocytes using tumor-derived metabolite lactate showed that the suppressive effects of lactate on macrophages extend beyond the tumor environment; such effects include induction of aerobic glycolysis, immunosuppression, and release of lactate (35). High lactate levels oppose lactate efflux from immune cells, leading to decreased cytokine production and cytotoxic activity (48); these effects have been observed in 2-AA tolerized macrophages (1, 10). 2-AA mediated tolerization increases glycolytic enzyme hexokinase-II and LDH levels, leading to lactate accumulation intracellularly and extracellularly. Intriguingly, because lactate can inhibit monocyte/macrophage differentiation (48), high lactate levels might hamper immune surveillance and promote host tolerance.

Reduced ATP levels in 2-AA tolerized immune cells may impact many cellular processes that require ATP, including cell signaling-related enzyme activity, histone acetylation, and cytokine production (36). Here, we showed that increasing ATP availability could reverse the effects in 2-AA tolerized cells, including restoring cytokine and H3K18ac levels in 2-AA tolerized cells, consistent with a prior study showing that lactate can blunt LPS effects on ATP production, thereby enabling inflammatory responses that require ATP to occur (49).

Regarding potential molecular mechanisms of immunomodulatory 2-AA, we noted that lactate dampened the activity of NF-κB, a major transcription factor that is important for inflammatory cytokine induction (1, 10). Lactate has been shown to suppress NF-κB activation (35) and downstream miR155 induction, thereby repressing inflammation (49). Here, we showed that accumulation of lactate, a byproduct of aerobic glycolysis, suppresses NF-κB signaling and proinflammatory cytokine production in 2-AA tolerized cells. The miR155 suppression that we observed could be one anti-inflammatory effect of lactate accumulation in 2-AA tolerized cells. Previously, miR155 has been shown to suppress inhibition of SOCS1 (suppressor of cytokine signaling-1) and to have an overall proinflammatory role in immune cell signaling (50). Together, these results suggest that lactate accumulation, whether endogenous or exogenous, suppresses inflammation.

In summary, the present data reveal that the 2-AA-mediated tolerization results from immunometabolic reprogramming, which is tightly linked to an interplay between metabolite biochemistry and immune networks. This reprogramming appears to lock cells in a deacetylation state. The presently revealed first-in-class metabolic reprogramming of immune cells by a QS signaling molecule offers a new avenue for designing and developing new therapeutics and interventions that can protect patients from recalcitrant persistent infections.

## Materials & Methods

### Ethics statement

The animal protocol was approved by the Institutional Animal-Care and Use Committee of Massachusetts General Hospital (2006N000093). No randomization or exclusion of data was applied.

### Tolerization assay

2-AA tolerization was performed as described previously (1, 12). Tolerized and non-tolerized mouse macrophage cells were supplemented with 10 mM sodium-D lactate (Sigma-Aldrich) and ATP (10, 20, or 50 µM, Sigma-Aldrich) for 3 h before 2-AA stimulation.

#### Pharmacological inhibitors

For the OXPHOS inhibition assay, non-pretreated and pretreated murine macrophage RAW264.7 cells were treated with UK5099 (10 µM, Sigma-Aldrich), etomoxir (2 µM, Sigma-Aldrich), and BPTES (10 µM, Sigma-Aldrich) 3 h prior to 2-AA stimulation. For glycolytic and lactate inhibition assays, non-pretreated and pretreated RAW264.7 cells were treated with 2-DG (1 mM, Sigma-Aldrich), sodium dichloroacetate (10 µM, Sigma-Aldrich), sodium oxamate (10 µM), and rotenone (1 µM, Sigma-Aldrich) 24 h prior to 2-AA stimulation.

### ATP determination

Extracellular ATP was detected with an ATP Assay kit (#ab83355, Abcam) according to the manufacturer’s instructions. Briefly, 50-µL standards and samples were added to a 96-well plate suitable for fluorescent analysis (black sides, clear bottom). A reaction mixture containing an ATP converter, probe, buffer, and developer mix was then added to all wells (50 µL) and incubated in the dark at room temperature for 30 min. Analysis was conducted at 535/587 nm.

### Lactate, acetyl-CoA, and pyruvate measurement

Lactate, acetyl-CoA, and pyruvate levels in culture media were determined with an L-Lactate Assay kit (ab65331, Abcam), a PicoProbe Acetyl-CoA kit (#ab87546, Abcam), and a Pyruvate Assay Kit (#ab65342, Abcam), respectively, according to the manufacturer’s instructions. Measurements were performed with at least three replicates.

### LDH and PDH activities

Culture media and cellular extracts were collected and centrifuged in preparation for LDH activity quantitation with a commercially available kit (#ab102526, Abcam). LDH activity was measured by a microplate reader (Tecan Infinite M200 Pro) at a wavelength of 450 nm in a kinetic mode at 25 °C for 30–60 min.

PDH activity was measured with an enzyme-linked immunosorbent assay (ELISA) kit following the manufacturer’s instructions (Abcam, #ab110671). Briefly, mt-protein extracts were incubated for 3 h at room temperature on plates pre-coated with anti-PDH antibody. Following incubation, plates were emptied, washed twice with stabilizer buffer, and incubated for 1 h at room temperature with detector antibody. PDH activity was determined by colorimetric measurement every 36 s at 450 nm and normalized to total protein.

### Statistical analysis

Whenever applicable, at least three independent experiments were performed. Error bars denote standard deviations (SDs). Statistical analyses were carried out using the GraphPad Prism software. One-way analyses of variance (ANOVAs) were conducted and followed by Tukey’s post hoc tests. Bacterial colony-forming unit (CFU) counts were analyzed using the Kruskal-Wallis non-parametric test with Dunn’s post-test.

## Acknowledgment

This work was supported by the NIH award R01AI134857, Shriner’s grant 85200, and The Massachusetts General Hospital Research Scholar Award to L.G.R. The Shriner’s grant 85132 to AAT and the Eleanor and Miles Shore Fellowship Program Award for Scholars in Medicine by Harvard Medical School to A.B. We thank Dr. Ramnik Xavier for critically reading the manuscript. The funders had no role in the study design, data collection, analysis, decision to publish, or manuscript preparation.

## Author Contributions

A.B. and L.G.R. conceived and designed the study. A.B., V.K.S, and A.C. performed experiments. A.B., V.K.S., and L.G.R. analyzed the data. A.A.T. contributed to the data interpretation. A.B and L.G.R. wrote the manuscript.

