## Supplementary Materials for "Immunometabolic hijacking of immune cells by a *Pseudomonas aeruginosa* quorum-sensing signal"

### Supplementary Information Text

#### Supplementary Materials and Methods:

#### Cell culture

All cells were maintained in 5% $CO_2$ at 37 °C. RAW264.7 cells carrying the NF-κB luciferase plasmid (IMGENEX, USA) and THP-1 Blue cells (human monocytic leukemia line with an NF-κB–inducible reporter (Invivogen) were maintained in Iscove's Modified Dulbecco's medium (IMDM, Invitrogen) and RPMI 1640 medium (Invitrogen), respectively. The media were supplemented with 10% heat-inactivated fetal bovine serum (endotoxin-free certified; Invitrogen), 2% penicillin/streptomycin, 2 mM L-glutamine, and 10 mM HEPES (all from Gibco). Cells were seeded in T-75 tissue culture flasks (Falcon, USA) and used between passages 2 and 3. For quality control, we tested the cells for mycoplasma with the PlasmoTest™ kit (Invivogen). Control siRNA and HDAC1-KD cells were grown in complete DMEM medium in presence of puromycine (10 µg/mL) (1).

#### Tolerization assay

RAW264.7 cells were plated (density of $10^6$/ml) in 6-well plates and grown overnight at 37 °C in 5% $CO_2$. Cells in treatment groups were pretreated with 800 µM 2-AA for 48 h; treated and non-treated cells were washed with 1X phosphate-buffered saline (PBS) and kept in fresh medium. Cells were stimulated with 400 µM 2-AA for the durations indicated in the figures. Similarly, THP-1 cells were tolerized (or not) with 400 µM 2-AA for 24 h and then stimulated with 200 µM 2-AA.

#### Cytosol and mitochondria fraction isolation

Cells ($1\times 10^6$) plated to incubate overnight were used for tolerization assays (described above). Mitochondria and cytosolic fractions of each group were isolated utilizing a Mitochondria Isolation Kit for Cultured Cells (#ab110170, Abcam) following the manufacturer's protocol. Following separation, a Bradford assay was conducted, cytosol and mitochondria fractions had 5~10 μg of protein. Samples were used for pyruvate assays.

**Immunoblot analysis**

Cells were plated at 6 × $10^5$ cells per well in six-well plates and then washed and lysed in RIPA lysis buffer containing 1 mM phenylmethylsulfonyl fluoride. Twenty-microgram samples of proteins were separated by electrophoresis (Kilo Dalton) in SDS-polyacrylamide gels; proteins were transferred to a 0.2-mm polyvinylidene fluoride membrane (Millipore, Billerica, USA) with a Bio-Rad semi-dry instrument. After blocking with 5% bovine serum albumin in Tris buffered saline containing 0.1% Tween-20 for 1 h at room temperature, the membranes were incubated with primary antibodies targeting ERR-α (ab76228, Abcam), MPC1 (D2L91, Cell signaling), pPDHE1-A type (Ser293) (ABS204, Milipore), GLUT1 (sc-377228), hexokinase-II (sc-130358), LDH-A (sc-137243), PDH (sc-377092), and anti-β-actin (sc-47778)(Santa Cruz Biotechnology) overnight at 4 °C. After washing, the membranes were incubated with anti-rabbit secondary antibody. Protein bands were detected by SuperSignal West Pico Chemiluminescent Substrate (Thermo Scientific) reaction, according to the manufacturer's instructions. The gels were visualized in a ChemiDOC Imaging system (Bio-Rad Laboratories, Inc., Hercules, CA). The bands were analyzed densitometrically with QuantityOne software (Bio-Rad).

**HDAC activity assay**

Nuclear fractions were used to measure HDAC activity with a non-isotopic assay that used a fluorescent derivative of epsilon-acetyl lysine (HDAC Fluorescent Activity Assay Kit, Active Motif), according to the manufacturer's instructions. Fluorescence intensity was measured using a Tecan plate reader (excitation wavelength 340–360 nm; emission wavelength 440–460 nm). HDAC activity was expressed in arbitrary fluorescent units.

**Luciferase assay**

After undergoing a 2-AA tolerization assay in a Luciferase Assay Kit (Promega)(2), RAW264.7 cells were washed with PBS and then lysed in kit-provided buffer.

**Quantification of global histone H3K18 acetylation**

Quantification of global lysine-specific histone acetylation was done with a Global Acetyl Histone K18 colorimetric assay kits (Abnova) according to the manufacturer's protocol.

**Measurement of TNF-α by ELISA**

TNF-α protein levels in culture supernatants of 2-AA tolerized and non-tolerized cells were measured by ELISA with Quantikine human/mouse TNF-α kits (R & D Systems) according to the manufacturer's instructions.

**miR155 assay**

We quantitated miR levels with a miR plate array (Signosis, Sunnyvale, CA) following the manufacturer's instructions. Briefly, 10–30 μg of extracted total RNA was utilized for hybridization in a 96-well plate pre-coated with miR oligo mix, including a pair of unique miR155 oligos that hybridize side-by-side with target miR, universal capture oligo, and biotin-labeled oligo. The streptavidin-HRP conjugate was used for miR155 detection. Chemiluminescence was determined for each well with a Tecan plate reader.

**Bacterial strains and growth conditions**

The PA14 strain of *P. aeruginosa*, also known as $Rif^R$ human clinical isolate UCBPP-PA14, was used (3, 4). The mutant strains Δ*mvfR* and Δ*pqsA* isogenic to UCBPP-PA14.The bacteria were grown at 37°C in Luria-Bertani (LB) broth under shaking and aeration or on LB agar plates containing appropriate antibiotics. PA14 cultures were grown in LB from a single colony to an optical density at 600 nm ($OD_{600}$) of 1.5 and diluted 1:50,000,000 in fresh LB media and grown overnight to an $OD_{600}$ of 3.

***In-vitro* mouse macrophage infection**

RAW264.7 mouse macrophage cells (1 x $10^6$) were plated in 6-well culture plates. Before infection, cells were washed in PBS and replenished with an antibiotic-free medium. The cells were infected with at a multiplicity of infection of 5 with *P. aeruginosa* [control (wild-type PA14) or an isogenic mutant (*mvfR* or polar *pqsA* mutant)] for 1 h. Supernatants were collected. Cells were washed three times and cellular extracts were prepared for biochemical analysis.

**Murine experiments**

A burn injury mouse model (1, 2, 5) was used to assess 2-AA effects on metabolites in 6-week-old CD1 male mice (Charles River Laboratories). Four days before burn and infection, mice were injected intravenously with 2-AA (6.75 mg/kg). Following administration of anesthesia, a full-thickness thermal burn involving 5–8% of the total body surface area was produced on the shaved mouse abdomen dermis and an inoculum of 1 × $10^4$ PA14 cells in 100 µL of $MgSO_4$ (10 mM) was injected intradermally into the burn eschar. CFU counts were assessed in 5 mice per group from muscle samples obtained from underneath the burn wound at 1 d, 5 d, and 10 d after burn and infection. Samples were homogenized in 1 mL of PBS, and then diluted and plated on LB-agar plates containing rifampicin (50 mg/L). We collected spleen samples from mice to perform metabolite analysis. Investigators were not blinded to experimental conditions and no randomization or exclusion of data points was applied. Spleen samples were collected from all the above mice 1 d, 5 d, and 10 d after burn and infection for ATP (#ab83355 Abcam), lactate (#ab65331, Abcam) and acetyl-CoA (#ab87546, Abcam) assays.

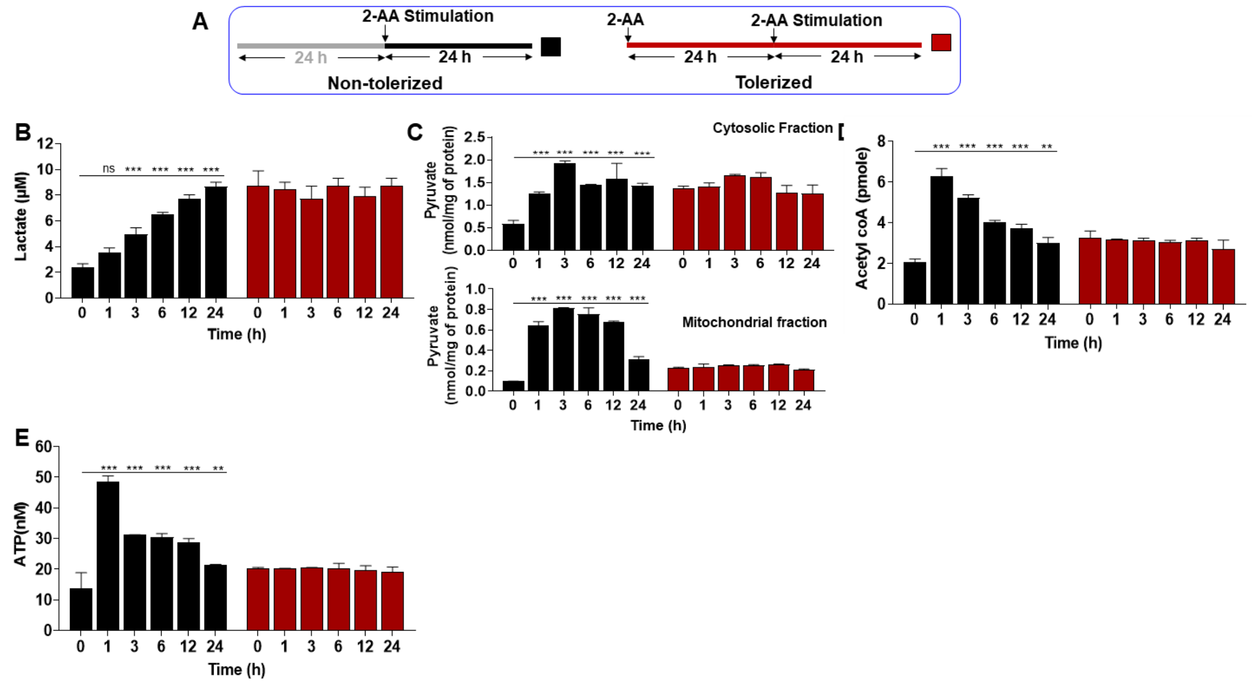


**Figure S1. 2-AA tolerization leads to decreased metabolites in human monocytes.** (A) THP-1 human monocytes were tolerized with 2-AA for 24 h or left untreated. In both cases, after 24 h, stimulation with 2-AA was conducted until the indicated times. Biochemical estimation of (B) intracellular lactate, (C) cytosolic- and mt-pyruvate, (D) acetyl-CoA, and (E) cellular ATP in non-tolerized and 2-AA-tolerized cells following 2-AA stimulation. Means ± SDs are shown (n = 3); ***p* < 0.01, ****p* < 0.001, and ns indicates no significant difference for one-way ANOVA.

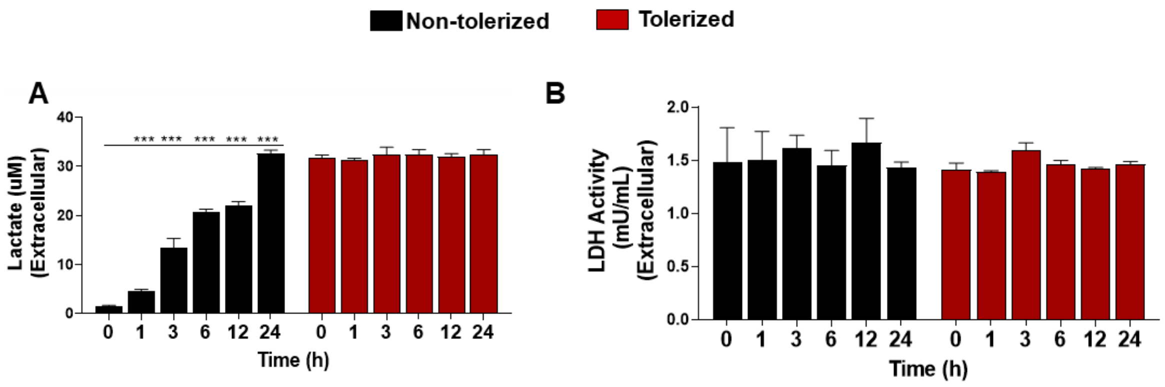


**Figure S2. Levels of extracellular lactate and LDH in 2-AA tolerized mouse macrophages.** (A) Extracellular lactate and (B) LDH activity were measured in 2-AA tolerized and stimulated cells. Means ± SDs are shown (n = 3); ***$p < 0.001$ for one-way ANOVA.

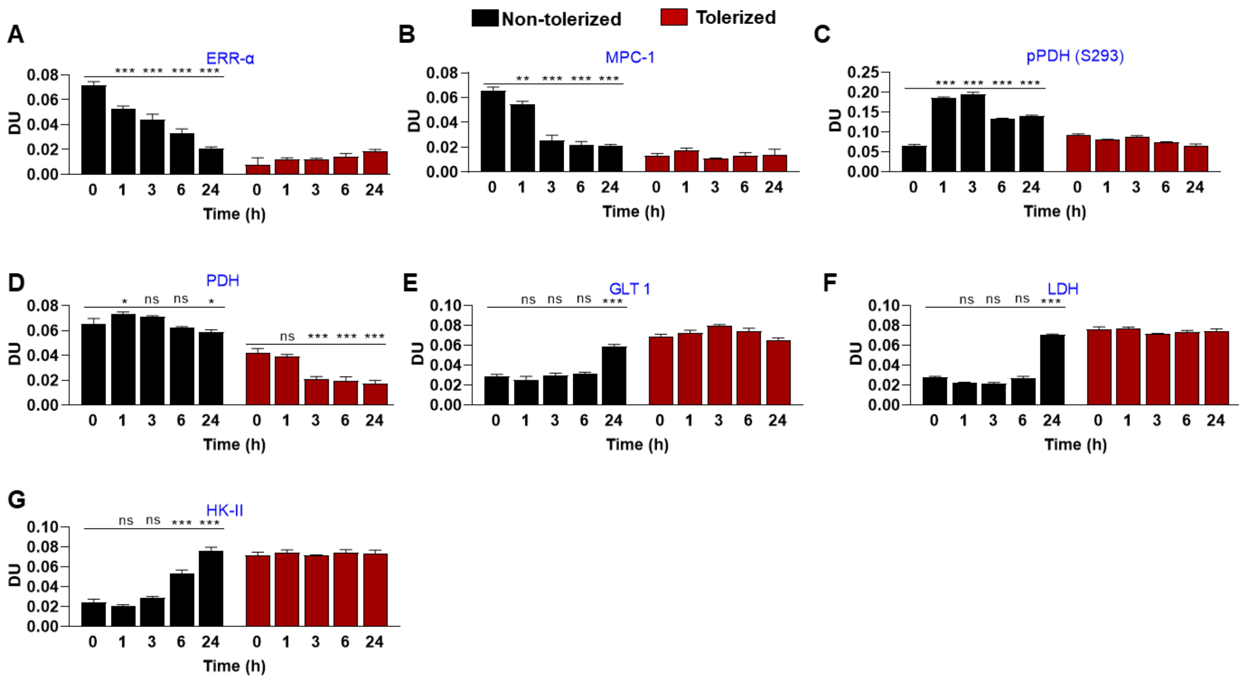


**Figure S3. 2-AA tolerization leads to increase in glycolytic flux and increase in lactate metabolizing enzymes in mouse macrophages.** (A-G) Histograms showing relative expression of ERR-α, MPC1, pPDH, PDH, Glut1, LDH, and HK-II proteins. Data are representative of three independent experiments. Means ± SDs are shown (n = 3); **p* < 0.05, ***p* < 0.01, ****p* < 0.001, and ns indicates no significant difference for one-way ANOVA. DU, densitometric unit.

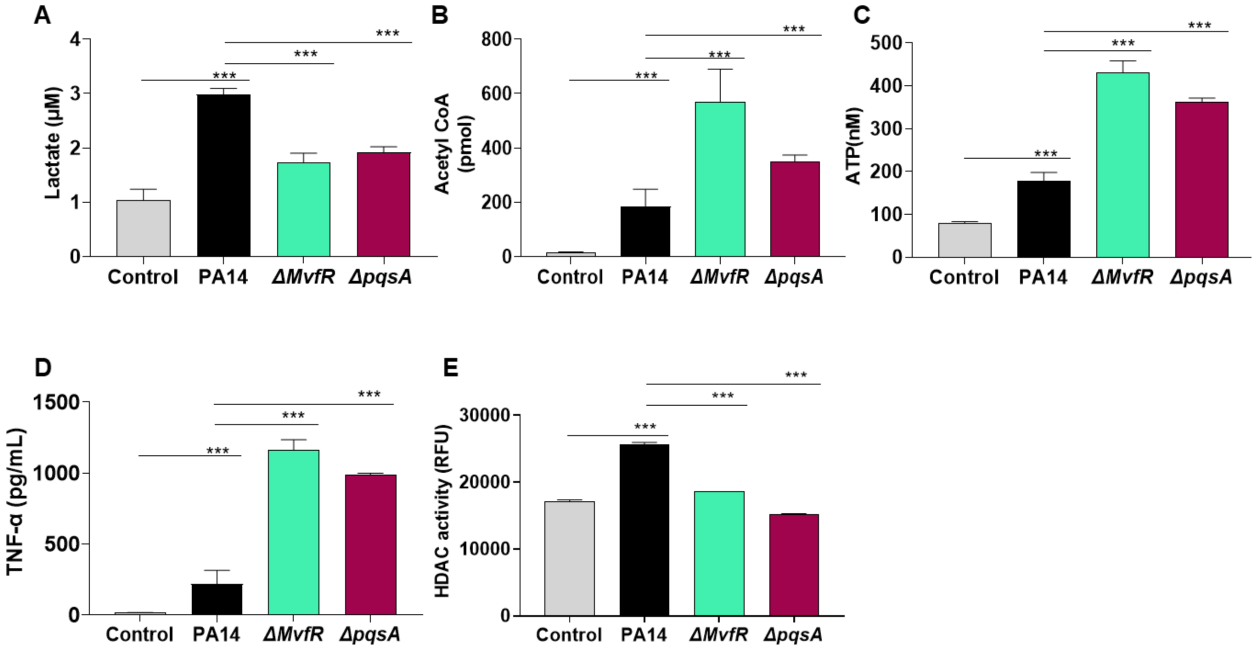


**Figure S4.** ***P. aeruginosa* regulates immunometabolic state of macrophages via MvfR-regulated 2-AA.** (A) Lactate, (B) acetyl-CoA, (C) ATP, (D) HDAC, and (E) TNF-α were measured in mouse macrophages after 1 h of 1:5 multiplication of infection with wild-type *P. aeruginosa* (PA14) and isogenic *mvfR* and *pqsA* mutants. Means ± SDs are shown (n = 3); ***$p < 0.001$ and ns indicates no significant difference for one-way ANOVA.
